# Multi-tasking Deep Network for Tinnitus Classification and Severity Prediction from Multimodal Structural Images

**DOI:** 10.1101/2022.05.07.491000

**Authors:** Chieh-Te Lin, Sanjay Ghosh, Leighton B. Hinkley, Corby L. Dale, Ana Souza, Jennifer H. Sabes, Christopher P. Hess, Meredith E. Adams, Steven W. Cheung, Srikantan S. Nagarajan

## Abstract

Subjective tinnitus is an auditory phantom perceptual disorder without an objective biomarker. Fast and efficient diagnostic tools will advance clinical practice by detecting or confirming the condition, tracking change in severity, and monitoring treatment response. Motivated by evidence of subtle anatomical or functional morphological information in magnetic resonance images (MRI) of the brain, we examined data-driven machine learning methods for joint tinnitus classification (tinnitus or no tinnitus) and tinnitus severity prediction. We propose a deep multi-task multi-modal framework for joint functionalities using structural MRI (sMRI) data. To leverage cross-information multimodal neuroimaging data, we integrated two modalities of 3-dimensional sMRI - T1 weighted (T1w) and T2 weighted (T2w) images. To explore the key components in the MR images that drove task performance, we segmented both T1w and T2w images into three different components - cerebrospinal fluid (CSF), grey matter (GM) and white matter (WM), and examined performance of each segmented image. Results demonstrate that our multimodal framework capitalizes on the information across both modalities (T1w and T2w) for the joint task of tinnitus classification and severity prediction. Our model outperforms existing learning-based and conventional methods in terms of accuracy, sensitivity, specificity, and negative predictive value.

## I. Introduction

Subjective tinnitus is an auditory phantom disorder characterized by the perception of internally generated elemental sounds, often described as ringing, humming, buzzing, chirping, or clicking, in the absence of externally identifiable sources. In its chronic phase, tinnitus manifests as a central nervous system disorder. Some prevailing hypotheses include maladaptive neuroplasticity, misappropriated attention, and dysfunctional striatal gating [1]–[3]. While hearing loss association is common, tinnitus severity or distress is often modulated by comorbid anxiety, depression, or mood disturbance. Bothersome tinnitus can degrade activities of daily life, disrupt sleep, and decrease work productivity [4]. There is a need for the scientific community to develop novel diagnostic methods to advance tinnitus management along its entire clinical course – starting from detecting or confirming the condition, progressing to tracking change in severity, and concluding in monitoring treatment response. This challenge in clinical tool development may be approached by machine learning methods [5] that categorize a patient into binary or multiple classes for tinnitus presence and use regression models of continuous clinical assessment values for estimating tinnitus severity.

Neuroanatomical and neurophysiological evidence point to tinnitus-related reshaping of the auditory pathway, with subtle reorganization of auditory areas (cortical and sub-cortical) over time [6], [7]. It has also been suggested that tinnitus is a neurodegenerative disorder which causes structural and functional changes in brain areas [8], [6], [9]. Researchers have explored various neuroimaging techniques to determine whether there are any structural abnormalities in the brain of tinnitus patients. White matter tracking using diffusion tensor imaging (DTI) shows alterations in both auditory and limbic areas [10]. Grey matter volume (GMV) extracted from structural MRI (sMRI) shows significant changes in various areas of tinnitus patients compared to healthy controls [11], [12]. A decline in GMV has been reported in ventromedial and dorsomedial prefrontal cortices, nucleus accumbens, anterior and posterior cingulate cortices, hippocampus and supramarginal gyrus [12]– [14].

In this work, our focus is to develop a data-driven framework for joint tinnitus classification into the presence or absence of auditory phantom percepts and estimation of tinnitus severity using regression models. Specifically, we aim to apply statistical and machine learning algorithms to sMRI data for the following objectives: 1) assess sMRI-based algorithm performance to differentiate tinnitus patients from healthy controls, and 2) identify the key sMRI features most strongly associated with clinical rating of tinnitus severity.

### A. Related Work

In this section, we briefly review prior work on neuroimaging-based methods for diagnosis of several neurological diseases and tinnitus. A critical consideration for optimizing clinical tool performance is choice of feature space for neuroimaging data extraction [15]–[17]. In terms of MRI feature representations, there are three categories: 1) voxel-based features [15] such as white matter (WM), gray matter (GM), and cerebrospinal fluid (CSF), 2) region-of-interest (ROI) based features [16] including regional cortical thickness, hippocampal volume, and GM volume, and 3) patch-based features [17]. Amongst these feature classes, voxel based features possess higher degrees of freedom - millions of voxels. These are independent of any hypothesis of brain structures. However, dimensional reduction of the high dimensional data remains an integral part of voxel based disease prognosis. Inspired by the tremendous success of deep learning [16], we consider deep features in contrary to hand-crafted feature.

In contrast to choosing a-priori features here we consider automated image based feature extraction using deep learning [16]. Many researchers have made promising contributions using deep learning methods for classification and regression models for symptom prediction in a variety of brain disorders, including Alzheimer’s disease [18]–[22], dementia [23], [24], and autism [18], [25]–[27] but none exists for tinnitus. A graph convolutional neural network that leverages both imaging and non-imaging information for brain analysis in large populations has been performed for autism [18]. Graph nodes are associated with imaging-based (functional MRI) feature vectors within the classification framework for autism and Alzheimer’s disease. A deep learning framework to classify Alzheimer’s disease (AD) based on whole brain sMRI using hierarchical structure of both voxel-level and region-level features has been proposed [19]. Yet another deep learning method based on sMRI has been introduced [20] to jointly detect AD disease and predicting clinical scores using structural MRI. Oh et al. [22] proposed a sMRI volumetric convolutional neural network (CNN) using an end-to-end learning approach using sMRI for AD classification and spatial attention visualization. Neuroimaging with computer-aided algorithms have made remarkable advances in dementia prognosis [23]. In particular, Xia et al. [24] proposed a novel dementia recognition framework based on deep belief network using functional MRI. Arya et al. [25] have developed a graph convolutional network by fusing structural and functional MRIs for autism classification. The aforementioned works motivate the current investigation using multimodal integration. Understanding the driving structural components of human brain towards capturing discriminating features for various brain disorders is of broader interest to computational neuroscience [28]. In particular, recent work on mild cognitive impairment (MCI) classification in [29] reported a preliminary studies on identifying the region-of-interests (ROI) with substantial influence in the classification task. We note that the study on correlation between GM and WM degeneration in various brain disorders is a potential experimental research direction [30]– [33]. Recent experimental findings suggest that tinnitus could potentially reorganize anatomical substrates in the brain [34], [35]. Motivated these works, we consider to explore how those anatomical substrates (GM, WM, and CSF) do substantially contribute our analysis framework.

We note that limited efforts using analytical and deep learning frameworks have been made for tinnitus detection and tinnitus severity prediction. Shoushtarian et al. [36] investigated the sensitivity of functional near-infrared spectroscopy (fNIRS) to differentiate individuals with and without tinnitus and to identify fNIRS features associated with subjective ratings of tinnitus severity. A machine learning method, including feature extraction and classification were applied to fNIRS data. An analytical approach based on whole-brain functional connectivity and network analysis was introduced for binary tinnitus classification [37]. A combined dynamic causal modeling and exponential ranking algorithm applied to EEG data yielded new insights into abnormal brain regions associated with tinnitus [38]. An unsupervised learning framework using a spiking neural network to analyze EEG data captured neural dynamic changes [39], extending earlier EEG based classification methods to capture neural dynamic changes [20]. To best of our knowledge, this is the first effort to explore deep learning for joint tinnitus classification and tinnitus severity prediction using structural MRI data.

#### Deep Multi-tasking in Medical Image Analysis

Multi-tasking networks are of special interest to the deep learning research community, where common discriminator features across multiple tasks could be learnt from input data [20], [40]– [44] including multimodal medical imaging data [45]. For instance, the multi-tasking networks in [40] combined models for regression and classification of lung nodules in CT images by stacking computational features derived from deep learning auto-encoder and CNN models with hand-crafted features were found to have superior performance to single task networks [40]. Similarly, authors in [43] introduced a novel multitasking deep framework for both regression and classification of the the Alzheimer’s Disease, wherein combining clinical data of various modalities (i.e., genetic information and brain scans) identified AD-relevant biomarkers. Multi-tasking of reconstruction and segmentation of brain MR images was successfully reported in [44] with impressive performance. Inspired by [44], we introduce a deep network framework for jointly solving classification models for tinnitus diagnosis and regression models for tinnitus severity prediction.

### B. Scientific Contributions

The main contributions are:

#### (a) Muti-tasking deep analysis of tinnitus disease

We propose a convolutional neural network built using a ResNet architecture for analysis of tinnitus using sMRI with whole brain 3D voxel-level features. We developed a multi-tasking deep framework for jointly performing tinnitus classification and tinnitus severity prediction. This includes a novel loss function for training a multi-tasking network. Our proposed method efficiently performs joint tasking of classification and clinical score prediction of tinnitus disease from the same set of deep features.

#### (b) Mutlimodal fusion

We attempt to intelligently combining features from both T1-weighted (T1w) and T2-weighted (T2w) sMRI within our network. Integration of multimodal structural imaging data within a joint framework leverages the strength of each modality and their inter-relationships for clinical tool performance.

#### (c) Structural controllability

Determination of imaging features (white matter, gray matter, cerebrospinal fluid) that contribute to tinnitus classification and tinnitus severity prediction performance by examining them individually and collectively.

#### (d) Novel multi-tasking loss function

We introduced a novel loss function for efficient learning in multi-tasking network. In particular, the proposed loss function jointly penalizes both classification and prediction by some form of convex combining of the individual losses. To mitigate the shortcomings of class imbalance in our dataset, we used focal loss [46] which focuses the training on the misclassification samples and largely bypasses the easy, correct classifications untouched. We also used data augmentation to avoid over-fitting during training. The loss which penalizes the regression is *L*_2_ error.

#### (e) Evaluation of clinical tool performance using the largest tinnitus dataset

We validated our proposed framework on the largest tinnitus dataset available to us from a collaborative research between the Univesity of California San Francisco and University of Minnesota. With elaborate experiments, we show that our method could achieve competitive classification accuracy over the state-of-the-art approaches. On the other hand, the proposed method offers satisfactory performance in predicting Tinnitus Functional Index (TFI) scores from the sMRI data of the subjects.

### C. Organization

The article is organized in the following manner. In Section II, we describe the dataset in detail and explicate the data processing pipeline. In Section III we explain our proposed deep learning method: fundamental architecture, fusion of sMRI T1w and T2w images, and dual-tasking framework. In Section IV, we present experimental results comprehensively and compare our method against the SVM benchmark. We present an elaborate discussion on our experimental finding, the connection and relevance of our work as compared to state-of-the-art methods for tinnitus prognosis. In Section VI, we provide the concluding comments.

## II. Methods

In this section, we introduce the sMRI datasets used in our analysis. We further discuss the details on data collection and preprocessing steps.

### A. Dataset

A total of 379 subjects provided the training dataset (T1w and T2w images) from two recruitment sites - University of California San Francisco (UCSF) and University of Minnesota. In total there were 183 tinnitus patients and 196 normal controls (Table I, top and middle rows show site distributions). As 44 UCSF subjects underwent two data acquisition sessions, data from the first or second session was chosen for use as the independent validation dataset (Table I, bottom row). Tinnitus or control binary labels and their corresponding structural images were fed into the model for the classification task. Further, there were 240 subjects from the training dataset also completed the Tinnitus Functional Index (TFI) [47], which measures tinnitus on a continuous scale from 0 to 100. Data from those subjects were fed into the regression model. There were 24 subjects from the independent validation dataset with TFI scores that were used for the further model evaluation.

**TABLE I.** Demographic and clinical information of the training/validation and independent dataset.

|  |  | Tinnitus | Control |
| --- | --- | --- | --- |
| UCSF | Number of subjects | 95 | 84 |
|  | Female/male | 33/62 | 41/43 |
| | Age (mean ± SEM) | 56.4 ± 13.6 | 44.3 ± 18.0 |
| University of Minnesota | Number of subjects | 88 | 112 |
|  | Female/male | 39/49 | 64/48 |
| | Age (mean ± SEM) | 50.5 ± 15.2 | 34.5 ± 13.8 |
| Independent Validation Dataset (UCSF) | Number of subjects | 24 | 20 |
|  | Female/male | 6/18 | 10/10 |
| | Age (mean ± SEM) | 57.3 ± 14.3 | 42.8 ± 19.1 |

### B. Data Processing

All T1w and T2w sMRI images are subjected to the same processing steps for quality control and image alignment. The clinical MRI scans contain irrelevant bony skull, soft tissue, and cervical regions, and variations in spatial orientation of acquired images that may affect clinical tool performance. Several steps for data co-registration are performed in order to align the images into the template space and exclude irrelevant or unnecessary regions. The FSL toolbox [48] is used for co-registration. Mask generation is the standard method, which has been proven to be reliable and consistent. The three steps are: 1) structural images are processed with a skullstrip step that removes the skull and other soft tissue regions that are not considered to be brain, 2) skullstripped images are registered into a MNI-152 2mm template space that aligns the structural data spatially, and 3) registered image data are processed with the intensity normalization step. These operations remove artifacts due to magnetic field inhomogeneity from different MRI scanners. These processed data are treated as the 3D volumetric input of the classification and regression task.

To identify and compare the features that drive performance in the volumetric whole-brain images and the brain region segmentations, we use the FSL toolbox [48], [49] to generate cerebrospinal fluid (CSF), grey matter (GM) and white matter (WM) regions. The segmentation process in Fig. 1 is applied to all available T1w and T2w images. We note that segmentation routine of FSL is based on a hidden Markov random field model that is optimized using the expectation-maximization algorithm. We take the skullstripped images (not containing skull and other soft tissue regions that are not considered to be in brain) as discussed above as the input for the segmentation routine. The pipeline takes into account the spatial information in terms of mutual information in local neighborhoods. It also includes a step of correcting the spatial intensity variations to overcome the bias field. Finally, the segmentation pipeline of FSL is able to classify the voxels of the structural MRIs of brain into white matter, gray matter and CSF. Note that we follow same form of FSL segmentation on both T1w and T2w images for both classification and regression tasks.

**Fig. 1.**
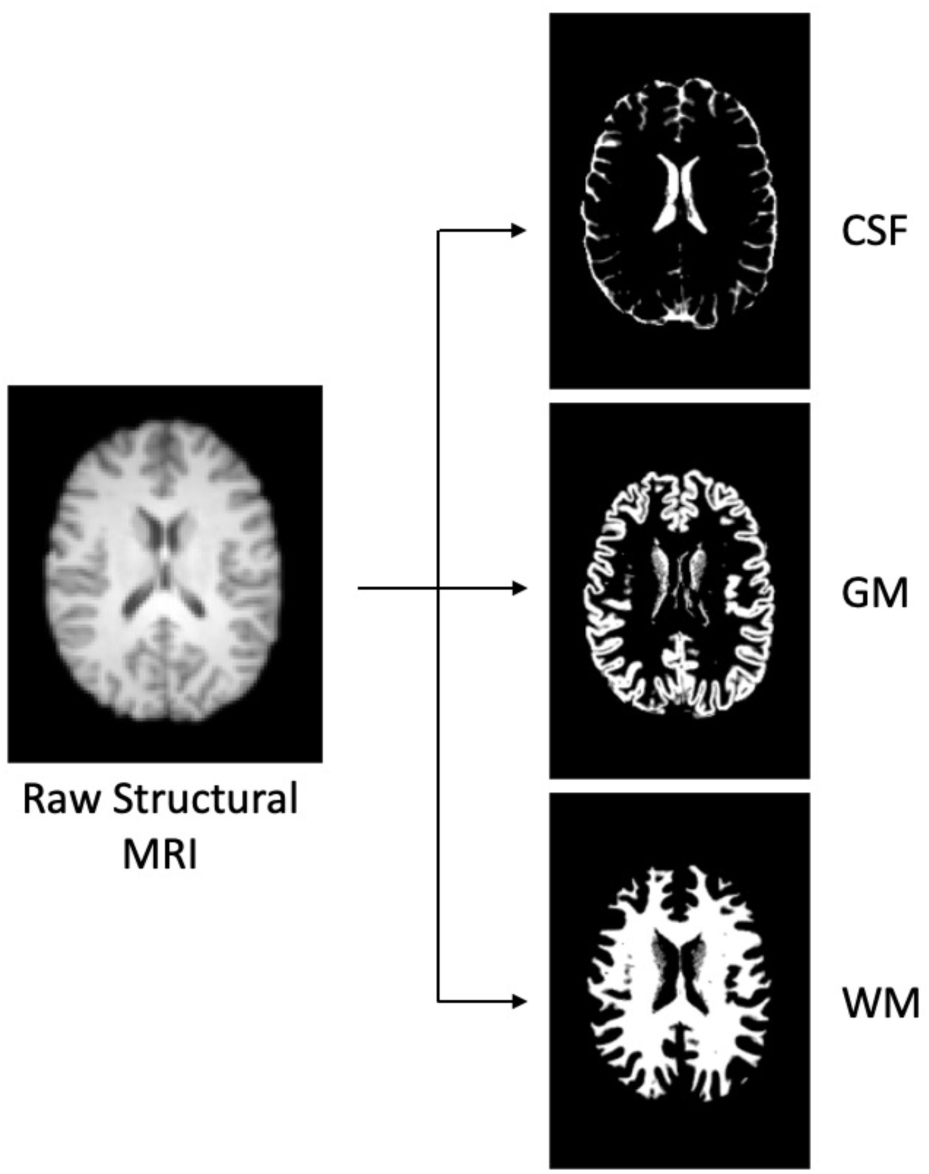
The workflow of brain segmentation process with FSL toolbox [49] to generate cerebrospinal fluid (CSF), grey matter (GM) and white matter (WM) segmentations.

## III. Algorithm

We start this section with the architecture of our proposed multitask algorithm. This is followed by a novel loss function to train the network for multi-tasking. Subsequently, we discuss the implementation details including data augmentation and transfer learning. The performance metrics to evaluate the performance of classification and TFI score prediction are also described. We also provide brief details of benchmark comparison algorithms which are compared later.

### A. Proposed Architecture

The algorithm for joint classification and regression tasks for tinnitus is built on ResNet-18 deep neural network architecture [42], which includes the residual information extracted from the previous layer and mitigates the adverse performance by using large number of layers. While we use the ResNet-18 architecture as our backbone, we then significantly modify it to incorporate both multiple modalities of imaging and the multi-tasking goals of our network. One attractive aspect of our method is the efficient integration of both modalities in structural MRI of each subject. In particular, the network consists of two parallel feature extraction sub-networks: one for T1w and another for T2w. The outputs from both sub-networks are concatenated. The fused composite features are further processed for joint tasking as shown in Fig. 2. Each of the sub-networks consists of 17 convolutional layers. The first convolutional layer takes the input and applies filters for the following convolutional layer input. Sixteen convolutional layers are wrapped into four convolutional blocks with four layers in each block. A detailed description of each convolutional block is displayed in Fig. 2. Note that same number of convolutional filters are used in each of convolutional blocks. For example, there are 64 filters in each convolutional layer of the first convolutional block. Each convolutional layer (within each convolutional block) follows with a batch normalization and a ReLu layer for more efficient gradient convergence (i.e. accelerate the training process) and only allows the positive values to pass to the next layer. Further, after each convolutional block, extracted features are pooled. The output from the fourth convolutional block of the two sub-networks are the features extracted from T1w and T2w data separately. We flatten each feature map into 1-D array to combine features. Next, we concatenate two 1-D arrays into a longer length of feature array. This simple fusion step essentially creates a double length fused feature array. The extended feature array is passed through two fully connected layers with different dimensions. The output layer of the final, fully connected layer FC-2 in Fig. 2(A) is used for multi-tasking: to simultaneously predict the class probability (via soft-max) and estimate the TFI score. The proposed deep learning method is referred to as multimodal multi-tasking convolutional network (MMCN).

**Fig. 2.**
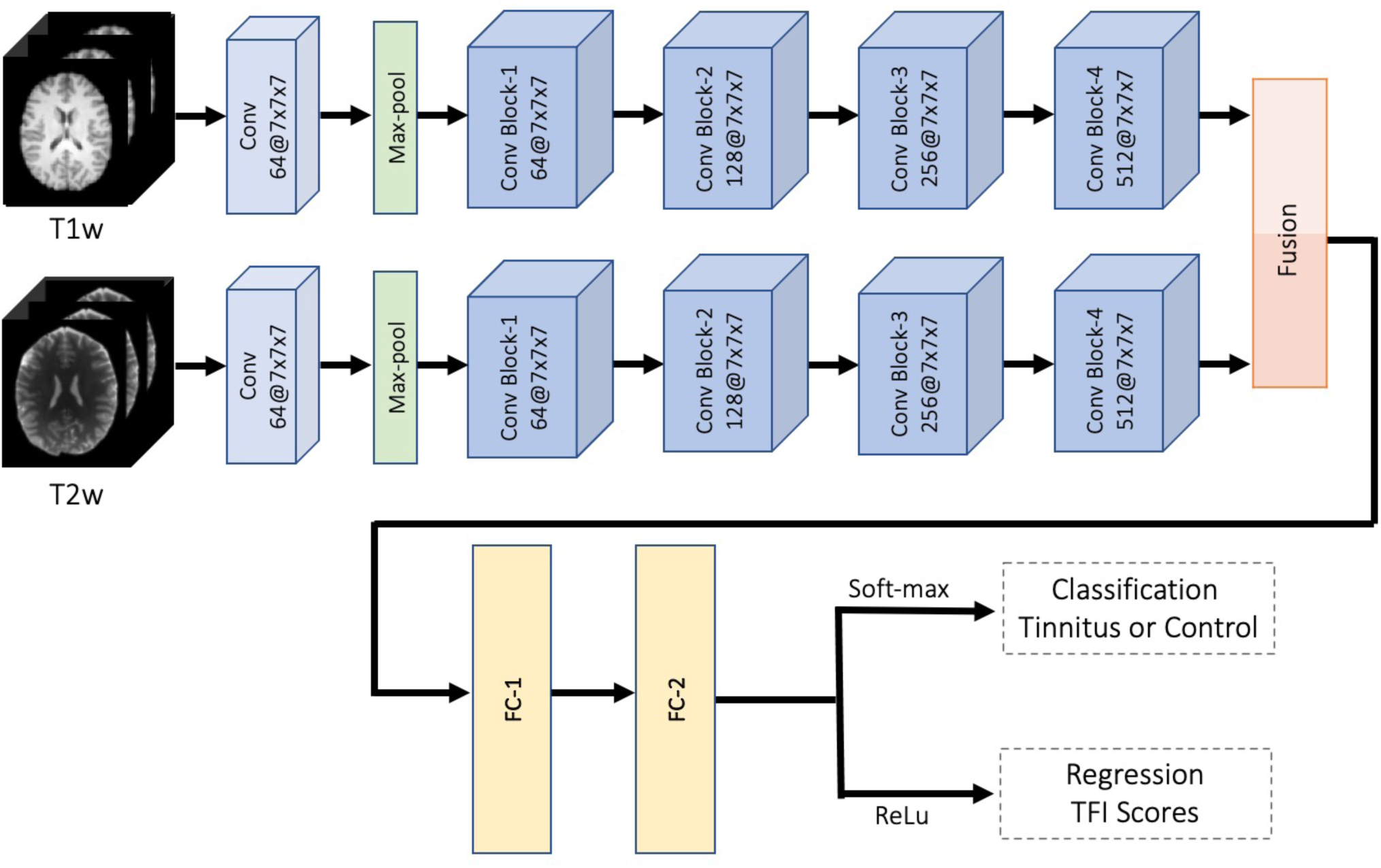
Overview of the proposed multimodal multi-tasking convolutional network (MMCN) for joint classification and regression of tinnitus. The pipeline takes the volumetric sMRI for both T1w and T2w as input in the first convolutional layer. There are four convolutional blocks in each sub-network. In each block, there are four convolutional layers that take the learned features from the previous block. Each convolutional layer in the block follows with a batch normalization and a ReLu layer to converge the gradient more efficiently and accelerate the training process. A pooling layer is applied at the end of each convolutional block to down sample and retain useful features. The filter size of each convolutional block are 64, 128, 256 and 512, respectively. A fusion step is applied after the final convolutional block to concatenate the features from each of the sub-networks. Further, two fully connected layers are used to learn non-linear combinations from the feature map. In the output layer, soft-max is used for tinnitus classification and ReLu is adapted for the regression task tinnitus severity prediction based on TFI scores.

### B. Proposed Multi-tasking Loss Function

#### 1) Loss Function for Classification

We experimented with two loss functions to optimize the classification performance, specifically, cross-entropy and focal loss [31]. Let the training set consisting of *N* number of subjects defined as 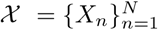. Each subject has a label 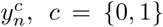, which indicates whether the subject is a healthy control or a tinnitus patient. In cross-entropy loss, the entropy *H(X)* is calculated for a random variable with a set of x in *X* discrete states and the corresponding probability *P(x)* as:

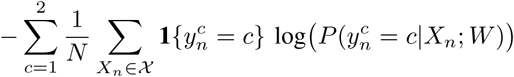

where **1**{*·*} is an indicator function; *W* is the collection of learned network coefficients and 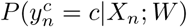 indicates the probability of subject *X*_*n*_ being correctly classified as the correct label 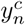. Note that **1**{*·*} = 1 if the condition {*·*} is true and 0 otherwise.

The labels of classification task in our setting are either tinnitus patient or healthy control. However, an imbalance dataset may lead to suboptimal training of the network. To deal with the limitation of class imbalance in the dataset, we further use focal loss, which is a modified version of the commonly used cross-entropy. Focal loss uses a modulating factor on top of the original cross-entropy equation, with tunable focusing parameter *λ ≥* 0. It is denoted as follows:

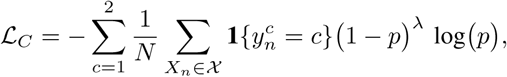

where 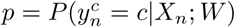. We empirically choose *λ* = 2. In fact, we experiment with different values of *λ*. Finally, we found *λ* = 2 to produce the most consistent results.

#### 2) Loss Function for Regression Task

To guide the training of regression task, we minimize *L*_2_ distance between the predicted and actual TFI scores. Suppose, there are *K* numbers of discrete TFI scores in our datasets such that 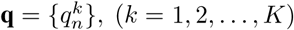. The mean squared loss between the estimated TFI score and the ground truth is

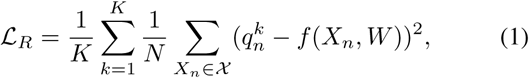

where *f* (*X*_*n*_, *W*) is the predicted TFI score of subject *n*.

#### 3) Loss Function for Multi-tasking

The training operation in the multi-task network is controlled by a loss function which could jointly penalize the error of both classification and regression tasks. In fact, we consider a convex combination of the cross-entry (or focal) and *L*_2_ loss. In particular, we use the following two composite losses:

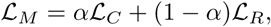

where *α* ∈ [0, 1] is a coefficient tuned using cross-validation. The convex combination controlled by *α* leverages an improved joint learning. The optimal value of *α* is set using cross-validation. We note that *α* could vary depending class imbalance and distribution of the ground-truth TFI scores in the datasets used. However, the protocal of setting *α* using cross-validation ensures the best possible training of the deep network.

### C. Implementation Details

The implementation of the proposed CNN model is based on the *Pytorch* library. The experiments were conducted on a NVIDIA GTX TITAN 16 GB GPU. We optimized the learning rate, and found the optimal value to be 10^*−*4^. We used a stochastic gradient descent (SGD) approach for optimization [50]. The network gradients while performing optimization are combined with the backpropagation algorithm. The learning rate for SGD are empirically set to 10^*−*3^.

We adopted the transfer learning to overcome the challenges with limited number of subjects. Transfer learning is widely used for obtained the weights for problems in the same domain to reduce the training time, to improve the overall performance, and to decrease overfitting. Here, we used pre-trained parameters from MedicalNet [51] as weights for all four convolutional blocks. Therefore, we only learnt the weights of the subsequent portions of the network. We followed the same form of transfer learning for both classification and regression prediction tasks. In summary, our proposed MCNN network learns the fully connected layers with transfer learning applied to the convolutional weights.

#### Data Augmentation

Availability of sufficient amount of data for adequately training a deep network is often a real concern in medical imaging. In fact, it is recommended to have at least 1000 samples of each class to train a classification model. To overcome the limited of this high data requirement, we performed data augmentation which also helps to avoid data over-fitting during the training. The goal of evaluating classification performance using an independent dataset was to validate the robustness of training.

We performed on-the-fly data augmentation using two strategies, i.e., i) randomly flipping the sMRI for each subject, and ii) randomly distorting the sMRI with non-linear transformation for each subject. The operation of randomly shifts introduces a reasonable perturbation to the training data for the network to learn the useful features. When combined with the first two operations, it could effectively augment the number and variability of available samples for training our MMCN model.

We examined the effectiveness of the proposed framework for the multi-modal multitask learning based test data and the independent dataset. To prevent the introduction of potential bias for not including the entire dataset, we constructed non-overlapping 5-fold cross validation in the training process. Specifically, we randomly selected 20% of the sample size from each class as the testing dataset, while the remaining 80% of the subjects were treated as the training dataset. This way we could utilize the existing available subjects to produce an unbiased performance.

### D. Performance Metrics

Classification performance was evaluated by five metrics, including classification accuracy (ACC), sensitivity (SENS), specificity (SPEC), positive predictive values (PPV), and negative predictive values (NPV). These standard metrics are defined as [19]:

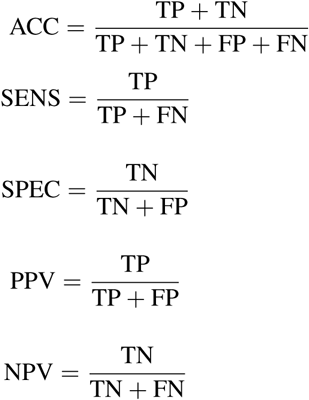

where TP, TN, FP, and FN denote the true positive, true negative, false positive, and false negative values respectively. For all of these five metrics, a higher value indicates a better classification performance.

Regression performance was evaluated using an r-squared (*r*^2^) metric, which is a statistical measure that represents the proportion of the variance for a dependent variable that’s explained by an independent variable or variables in a regression model [52]. The definition of the coefficient of determination is as follows. Suppose the ground-truth and predicted TFI scores of n-th subject are given by *q*_*n*_ and 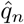. There are total *N* subjects used for testing to evaluate the regression performance. Also, we refer the mean of the ground-truth scores of all subjects as 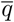. Then, the r-squared (*r*^2^) metric is defined as:

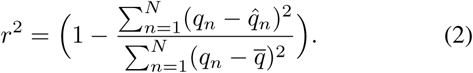

We clearly see from the above expression in (2) that it will take values between 0 and 1. At perfect prediction of TFI scores, by setting 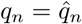 for all *n* in (2), we obtain *r*^2^ = 1. In summary, a higher value of *r*^2^ indicates a better regression performance achieved by the proposed method.

### E. Influence of Structural Regions on Multi-tasking

We perform detailed study to understand whether it is the whole brain MRI or a particular structural regions such as CSF, WM or GM that plays a dominant role in the multimodal multi-tasking performance of our method. In particular, both T1w and T2w images are first segmented into respective GM, WM, and CSF components. Thus, including whole brain (WB), we got total 4 sets of data for each subject. The detailed steps of using WB data into our network are explained in Fig. 2. For the analysis using these three structural segments, we simply substitute the respective segment of both T1w and T2w images. For example, to perform the joint tasking based on gray matter (GM), we use GMs derived from T1w and T2w respectively as inputs. We follow simlar steps for both WM and CSF based joint tasking. Note that we perform the training and test for these four cases completely independently. Therefore, the independent way of training the network allows to learn the joint tasking selection parameter *α* at best possible fraction. However, we use same form of transfer learning and data augmentation for all four cases.

### F. Benchmark Methods

The proposed MMCN method was first compared against two conventional learning-based methods - least squares support vector machine (LS-SVM) [53] and K-nearest neighbor (KNN) [54]. Beyond this, MMCN was compared with a state-of-the-art deep-learning method, the hierarchical fully convolutional network (H-FCN) [19], which has been used to extract useful features from imaging data to classify Alzheimer’s disease. To the best of our knowledge, there is no existing deep learning methods for tinnitus classification or severity prediction. We now briefly summarize the three benchmark methods.

1. LS-SVM: We use a modified version of support vector machine (SVM), a widely used classification model in neuroimaging analysis [53], [55]. We directly train a least squares SVM model on T1w whole brain volumetric MRI images that contains all available structural information. The trained model is applied on the test data to obtain final classification results in the test sample.
2. KNN: K-nearest neighbor is another popular, classical method used for performing unsupervised classification. This simple machine learning algorithm is based on the distance between feature vectors [54]. The k-NN algorithm classifies new unknown data points by finding the most common class among the k-closest centroids. In our case, T1w whole brain volumetric MRI of each subject is treated as sample points for k-NN algorithm. Test subjects are classified based on neighborhood of the learned centers (k-means).
3. H-FCN: H-FCN was a recent deep learning network with a unified discriminative feature extraction algorithm for successful classification of Alzheimer’s disease (AD) using volumetric 3D sMRI data [19]. The hierarchical fully convolutional network (H-FCN) method uses the same data format as our method, MMCN, and motivates a comparison.

## IV. Results

### A. Classification Performance

In this section, we focus on classification performance of the proposed MMCN deep model and compare performance against contemporary methods. We focus on three aspects: 1) exhibit performance of our proposed MCNN pipeline with different segmented regions, 2) compare performance against benchmarks, and 3) evaluate the classification accuracy on an independent dataset.

Tinnitus classification results for whole-brain and segmented brain regions, specifically CSF, GM and WM, are summarized in Table II. For all metrics, higher values indicate better performance. The bolded best performance metric highlights the corresponding structural region. Overall the whole brain outperforms the other three segmented brain regions. In particular, it achieves 72.9% in accuracy, 70.8% in sensitivity, 75.4% in specificity, 69.7% in positive predictive value, and 75.1% in negative predictive value, and the highest r2 value. Gray matter sMRI input has best performance metrics in sensitivity (77.7%) and positive predictive value (70.6%). While whole brain sMRI images provide the most useful features to drive the performance, other brain regions retain partial information.

**TABLE II.** Influence of structural regions of brain on multi-tasking performance using our multimodal method.

| Structural Regions | ACC (%) | SENS (%) | SPEC (%) | PPV (%) | NPV (%) | $r^2$ |
| --- | --- | --- | --- | --- | --- | --- |
| Whole Brain (WB) | **72.9 ± 4.2** | 70.8 ± 1.8 | **75.4 ± 2.1** | 69.7 ± 3.7 | **75.1 ± 3.9** | **0.429** |
| Cerebrospinal Fluid (CSF) | 71.2 ± 4.0 | 73.7 ± 4.2 | 64.3 ± 2.8 | 66.9 ± 5.1 | 74.7 ± 1.6 | 0.235 |
| Gray Matter (GM) | 72.0 ± 2.5 | **77.7 ± 2.0** | 66.9 ± 4.1 | **70.6 ± 4.2** | 73.7 ± 3.6 | 0.380 |
| White Matter (WM) | 70.1 ± 3.9 | 72.1 ± 3.5 | 68.1 ± 3.0 | 67.5 ± 2.8 | 74.6 ± 2.6 | 0.356 |

In Figure 3, we show the classification performance on the independent dataset. Note that our proposed MMCN method achieves around 70% accuracy using whole brain 3D data (both T1w and T2w). Here whole brain data refers to the case where no segmentation is performed on the 3D sMRI images. To further investigate the impact of segmented brain regions, we report the results using CSF, GM, and WM. The bar plots in Fig. 3 indicates that among these three brain regions, WM offers superior performance with respect to accuracy, sensitivity and specificity. WM segmented from sMRI images captures more prominent and representative features of tinnitus.

**Fig. 3.**
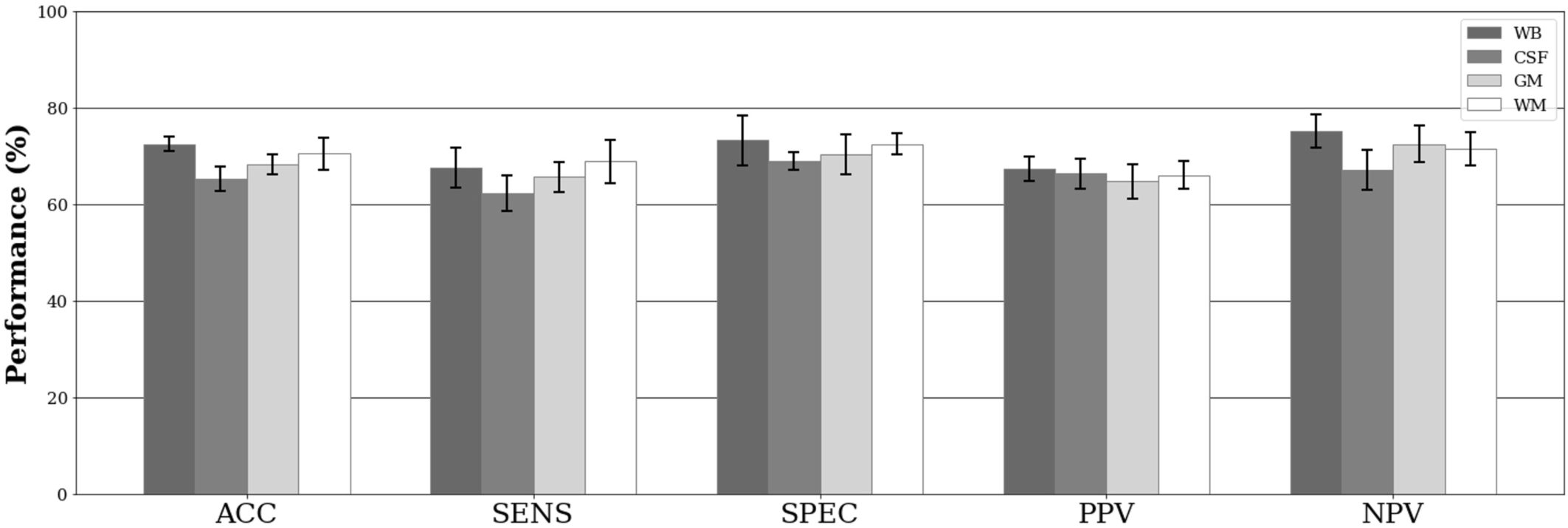
The MCNN performance on UCSF independent dataset.

Taken together, the proposed MMCN method offers best performance results using the whole brain in terms of all metrics except SENS and NPV. That said, GM achieves superior performance in terms of SENS and NPV. Based on this experiment, we conclude that GM contains tinnitus descriptive features at best among three structural regions. Whole brain 3D structural image data for input to MMCN appears to be the single best choice.

### B. Regression Performance

The task in regression modelling is to predict tinnitus severity based on TFI scores from the sMRI data. In Figure 4, we provide the scatter plots of whole brain, and the other three brain segmented regions, CSF, GM and WM, with corresponding prediction versus ground truth values. The x-axis shows the actual TFI scores (0 to 80) and the y-axis shows the predicted TFI scores (0 to 100). The shaded area represents the 95% confidence interval of the corresponding linear approximation. The slope in each scatter plot reveals whether the regression exhibits a positive or negative trend. The evidence of positive slope in each case validates our method can predict TFI scores reasonably well from sMRI images. The *r*^2^ value is the percentage of the variance in the dependent variable that the independent variable can explain. Whole brain outperforms the other three segmented regions with *r*^2^ = 0.429. This resonates with superior whole brain classification performance.

**Fig. 4.**
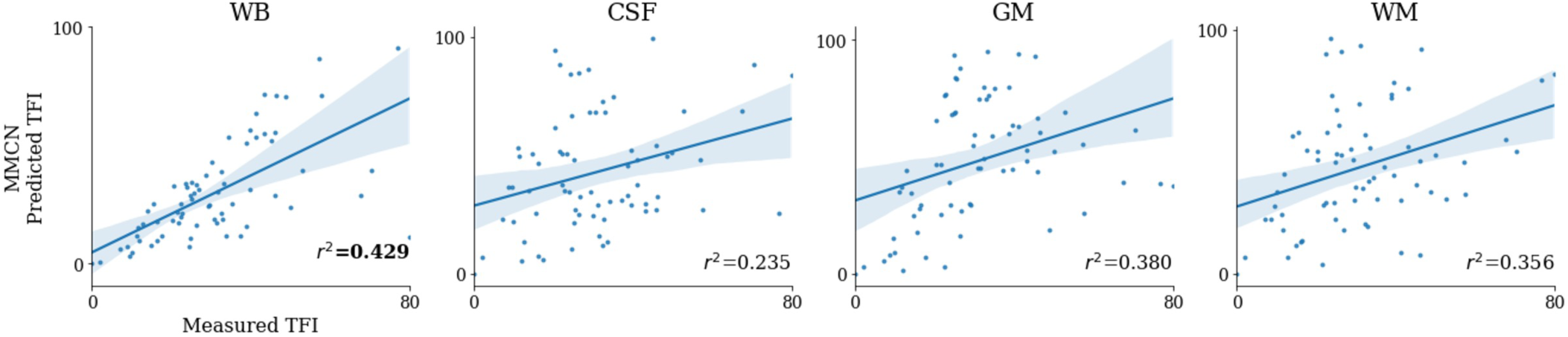
Scatter plots of measured TFI scores with predicted TFI scores.

### C. Benchmark Comparisons

We compare our MMCN method with two classical machine learning methods and one state-of-the-art deep learning approach in Table III. Note that our method is capable of performing both tasks: tinnitus classification and tinnitus severity. In addition, it is a multimodal methods approach, handling two modalities of sMRI data as input in parallel. In contrast, the three existing approaches are unimodal - they are capable of classification based on only one modality at a time, either T1w or T2w.

**TABLE III.** Comparison of classification performance. Note that all the existing methods can support only a single modality. For these methods we report the best performance using either T1w or T2w. Our proposed approach MMCN takes both both T1w and T2w as inputs.

|  | ACC (%) | SENS (%) | SPEC (%) | PPV (%) | NPV (%) |
| --- | --- | --- | --- | --- | --- |
| LS-SVM | 56.3 ± 3.1 | 57.8 ± 2.1 | 53.1 ± 2.1 | 57.3 ± 1.3 | 53.2 ± 3.4 |
| KNN | 53.2 ± 2.6 | 52.4 ± 2.9 | 55.2 ± 2.5 | 50.7 ± 1.9 | 58.1 ± 1.1 |
| H-FCN | 67.6 ± 3.4 | 70.1 ± 4.2 | 65.0 ± 2.9 | 69.2 ± 3.6 | 62.3 ± 4.5 |
| MMCN | **72.9 ± 4.2** | **70.8 ± 1.8** | **75.4 ± 2.1** | **69.7 ± 3.7** | **75.1 ± 3.9** |

MMCN has superior classification performance compared to the best performance of these three methods. LS-SVM and KNN produce best performance using T1w images; whereas H-FCN produces best performance using T2w images. Notice that our MMCN method outperforms both classical machine learning methods by a large margin across all five metrics (Table III). In direct comparison to the state-of-the-art H-FCN method, our MMCN method outperforms in all five metrics. The proposed MMCN achieves superior state-of-the-art classification performance. Two key factors may be contributing to superior performance. First, the proposed MMCN method jointly learns the discriminative features of structural MRI along with the classifier and regressor, and thus those learned features can be more suitable for subsequent classifiers/regressors. The proposed deep architecture perhaps can capture the discriminative features from the samples more accurately than H-FCN. Second, MMCN explicitly integrates both T1w and T2w modalities of sMRI data. Our multimodal deep network MMCN efficiently exploits cross-information present in both modalities.

## V. Discussion

Deep convolutional neural network (CNN) methods have achieved extraordinary success in medical image alaysis by extracting and adapting the highly discrimative features present in the images. One key research focus in deep learning based image analysis is on further improving the classification accuracy by applying insightful architectures and modules. In this work, we introduce a novel deep learning framework for classification of tinnitus subjects from its structural MRI data. Besides the intuitive (binary) classification result, the method can also output TFI scores as an indicator of the severity of the disease. From mathemitical point of view, this deep module provides a data-driven nonlinear relationship between MRI volume (consists of thousands of voxels) and TFI score. A remarkable aspect of the proposed network is that both classification and score prediction are achieved by same set of learnt deep features and nonlinear weights. Finally, our proposed multi-task deep model could be an efficient tool for structural MRI analysis to determine whether a patient is having tinnitus or not and if so, also to predict the severity of the disorder. Thus, we provide a fast and efficient diagnostic tool is essential early detection, monitoring clinical trails and tracking the progression of tinnitus.

Considering the lack of interpretability for CNN-extracted features, it is difficult to directly connect the classification/prediction results with the morphological attributes of MRI data. To improve the interpretability of our deep network module, we made an attempt by segmenting the MRI data into three micro-structural components - gray matters, white matters and cerebrospinal fluid (CSF). Then, we studied the multi-tasking performance by using each of these micro-structural components separately and compare them with the results obtained from unsegmented (whole brain) data.

One important aspect of our proposed method is the integration of multimodal structural MRI data within out joint framework to capitalize on the strength of both T1w and T2w modalities. The multimodal fusion of deep features allowed to exploit cross-modal information. Finally, we are able to achieve superior performance rather than the one obtained from each modality separately. By maximizing joint information available in both modalities, we integrate the learned features within our pipeline. We note that none of the methods compared in Section IV can support multiple modality for classification. We also add that our proposed method does not require much of preprocessing of the input data, unlike multi-tasking scheme in [20], which includes computationally intensive step of landmark patch extraction.

## VI. Conclusion

In this paper, we introduced a novel deep learning framework for simultaneous tinnitus classification and tinnitus severity prediction using structural MR imaging data. In particular, we integrated deep features from of two modalities - T1w and T2w of the available MRI data. Experiment results confirmed that the proposed method MMCN outperforms several recent methods for both tinnitus classification. In future, the proposed framework may potentially be deployed in real-time health care settings to confirm tinnitus, track change in severity, and monitor treatment response.

## Notes

### Competing Interest Statement

The authors have declared no competing interest.

